# Viscoelasticity of single macromolecules using Atomic Force Microscopy

**DOI:** 10.1101/2020.05.21.107888

**Authors:** Shatruhan Singh Rajput, Surya Pratap S Deopa, Jyoti Yadav, Vikhyaat Ahlawat, Saurabh Talele, Shivprasad Patil

**Affiliations:** Department of Physics, Indian Institute of Science Education and Research Pune, Dr. Homi Bhabha Road, Pashan, Pune 411008, India; Department of Chemistry, Indian Institute of Science Education and Research Pune, Dr. Homi Bhabha Road, Pashan, Pune 411008, India

**Author notes:** SP designed the research and experiments to be performed. SR and SD carried out all the experiments, performed the calculations and analyzed the data. VA helped in experiments and ST helped in developing the interferometer based AFM. SP, SR and SD wrote the article. JY expressed and purified the protein.

**Keywords:** Single molecules, viscoelasticity, Atomic Force Microscope

## Abstract

We measured viscoelasticity of single protein molecules using two types of Atomic Force Microscopes (AFM) which employ different detection schemes to measure the cantilever response. We used a commonly available deflection detection scheme in commercial AFMs which measures cantilever bending and a fibre-interferometer based home-built AFM which measures cantilever displacement. For both methods, the dissipation coefficient of a single macromolecule is immeasurably low. The upper bound on the dissipation coefficient is 5 × 10^−7^ kg/s whereas the entropic stiffness of single unfolded domains of protein measured using both methods is in the range of 10 mN/m. We show that in a conventional deflection detection measurement, the phase of bending signal can be a primary source of artefacts in the dissipation estimates. It is recognized that the measurement of cantilever displacement, which does not have phase lag due to hydrodynamics of the cantilever, is better suited for ensuring artefact-free measurement of viscoelasticty compared to the measurement of the cantilever bending. We confirmed that the dissipation coefficient in single macromolecules is below the detection limit of AFM by measuring dissipation in water layers confined between the tip and the substrate using similar experimental parameters. Further, we experimentally determined the limits in which the simple point-mass approximation of the cantilever works in off-resonance operation.

**Significance Statement:** Single Macromolecules, including unfolded proteins bear rubber-like entropic elasticity and internal friction characterized by finite dissipation coefficient. Direct measurement of this viscoelastic response is important since it plays a significant role, both in polymer physics as well as protein folding dynamics. The viscoelastic response of single polymer chain is difficult and prone to artefacts owing to the complications of hydrodynamics of macroscopic probe itself in the liquid environment. Using a special atomic force microscope, which allows quantitative estimate of viscoelasticity in liquid environments, we measured viscoelastic response of single molecule of Titin. We report here that the dissipation coefficient is below the detection limit of our experiments - with upper bound which is less than reported values in the literature.

---

The viscoelasticity of single proteins and other biologically relevant macromolecules is essential to understand how they function in single molecule limit. Atomic Force Microscopy is used to measure viscoelasticity of single macromolecules and other nano-scale systems owing to its unprecedented spatial resolution in physiological conditions (1–11). In typical AFM experiment, mica or Au substrate is sparsely coated with the biological macromolecule and is placed in the liquid cell. An oscillating sharp probe attached to a cantilever spring is then brought close to a molecule. The protein is allowed to attach to it from either C or N terminus through nonspecific binding. The bending in the cantilever beam as well as the phase and amplitude of cantilever oscillations is measured as the molecule is pulled away from the surface. The cantilever bending provides the amount of force applied on the protein as it is slowly stretched. The amplitude and phase may provide the viscoelastic response of the molecule at different extensions (7, 10, 11).

A solution to an appropriate equation of motion for the cantilever whose tip is tethered with protein provides a relationship between measured parameters (amplitude and phase) and molecule’s viscoelastic properties, namely the stiffness and dissipation coefficient (7, 12). This approach has been extremely successful for experiments performed in vacuum or air (13–15), however quantification of viscoelastic response from observed quantities is not straightforward in liquid medium. This is due to complications of cantilever dynamics in liquids and the method employed to excite cantilever in a liquid cell. Since the measured cantilever response is mixture of hydrodynamic forces and the forces due to stretched molecule, the quantification and separation of these two is essential to accurately determine the viscoelasticity of single molecules (5, 7, 8, 16–18). It has been recognized that the hydrodynamic forces play a crucial role in determining the phase of the tip motion without the molecule. This phase lag becomes important in accurately determining viscoelasticity of the molecule (5, 6, 12, 19–22). There are theoretical works which provide expressions to predict the phase of a cantilever oscillating in liquid environments (7, 12, 22). It has also been suggested that the use of phase and amplitude to determine the viscoelastic response of nanoscale systems in liquids leads to artefacts (7). Moreover, it has been noticed that researchers seldom find the phase lag predicted by theoretical models in experiments. This can be attributed to the phase contributions coming from variety of unknown sources. These sources are responsible for the randomness of experimentally observed phase once the cantilever is immersed in liquid and is yet to be tethered with molecule. It is extremely difficult to account for such contributions in theoretical models. Therefore it becomes important to make sure there are no contributions from the extraneous sources before the measurement is performed. We refer to phase contributions from such sources as extraneous phase.

In this work, we ensured that extraneous contributions to the phase are not present and performed dynamic, off-xresonance force spectroscopy experiments on Titin I27_8_ to measure its viscoelasticity. The measurements are performed with both, a conventional deflection-detection type AFM and a fibre-interferometer based home-built AFM. The fibre interferometer measures cantilever displacement and the deflection detection scheme measures the cantilever bending at it’s free end. The later is widely used in commercial AFMs. It is pointed out recently that interpreting phase of the cantilever bending as dissipative signal is inaccurate owing to altered boundary conditions due to molecule’s viscoelasticity. Using the solution to the Euler-Bernoulli equation of the rectangular cantilever beam, it is proposed that the X and Y component of the oscillations in cantilever bending exclusively determine the stiffness and dissipation coefficient respectively (7). We show here that even the use of X and Y components to estimate stiffness and dissipation coefficient are not free from artefacts in the presence of extraneous phase and results in cross-talk between the stiffness and dissipation channels. Such artefacts also appeared when the measurements are not strictly off-resonance, a major concern when operating in viscous media.

The theoretical models predict no phase lag due to hydrodynamics for the displacement of the cantilever’s free end in viscous medium. Therefore, in interferometer based detection, it is easy to ensure that extraneous phase due to sources other than hydrodynamics is not present. The bending of the cantilever, however has a finite phase lag due to hydrodynamics and it is difficult to separate the phase due to hydrodynamics and the extraneous phase. After ensuring that there are no contribution to the phase from extraneous sources, we compare the stiffness estimates from measurements of the displacement and the bending of cantilever’s tip end.

Using both, the interferometer based AFM and the deflection detection type AFM, we found that the dissipation in unfolded Titin I27_8_ octomer is immeasurably low. The interferometer based AFM, however clearly show evidence of dissipation in water layers from Y-component of the displacement oscillations. Since the dissipation is detected in water layers and not in unfolded protein molecule using the same instrument, we conclude that the dissipation in single macromolecules is indeed immeasurably low. The estimated detection limit of our instrument and hence the upper bound on the dissipation in single unfolded protein is 5 × 10^−7^ *kg/s*. We compared the stiffness estimated from the experimental data from both detection schemes. They match very well and also with stiffness using static method (derivative of force). The stiffness of unfolded I27_8_ is in the range of 10 mN/m.

For measurement of viscoelasticity at nanoscale, two different approaches are debated for accurate analysis of the experimental data. In one approach the viscoelastic response is included in equation of motion describing the cantilever dynamics and in another the viscoelasticity is included in the boundary conditions. In the first approach the dynamics is reduced to the first eigen mode of the cantilever beam and the cantilever is treated as point-mass. Recently, there are questions raised over the appropriateness of the point-mass model (7). Here, We clearly demonstrate that the point-mass model is adequate in case of displacement measurements performed using interferometer based AFM. We show that within certain limits of change in cantilever amplitude due to viscoelastic response of the molecule, both these approaches give identical results.

## Materials and Methods

### AFM

We used two types of AFMs to perform measurements. For deflection detection measurements, we used commercial AFM (JPK Nanowizard II, Berlin, Germany) in which a narrow laser beam incident at the end of the cantilever gets reflected to a four quadrant, position sensitive photo-detector. The position of the reflected beam on the detector corresponds to angle *dy/dx* due to the bending in the cantilever (see Fig. 1(a)). In another home-built AFM, a fibre-based interferometer is used (Fig. 1(b)). In brief, a partially coated end of the optical-fibre is aligned parallel to the back of a cantilever-end with the help of a nano-positioner. Part of the Laser (1380 nm) is reflected from this semi-mirror at the end of the cantilever. The rest of the light falls on the back of the cantilever and reflects back into the fibre. These two reflected lights interfere to form an interference pattern on a photo-detector. For more details about the instrument see reference (23). The signal from this photo-detector corresponds to cantilever displacement (y) as opposed to angle (*dy/dx*) at the free end due to the cantilever bending. This is a crucial difference, since the solution used to quantify stiffness and dissipation is different for angle *dy/dx* and for displacement *y*.

**Fig. 1.**
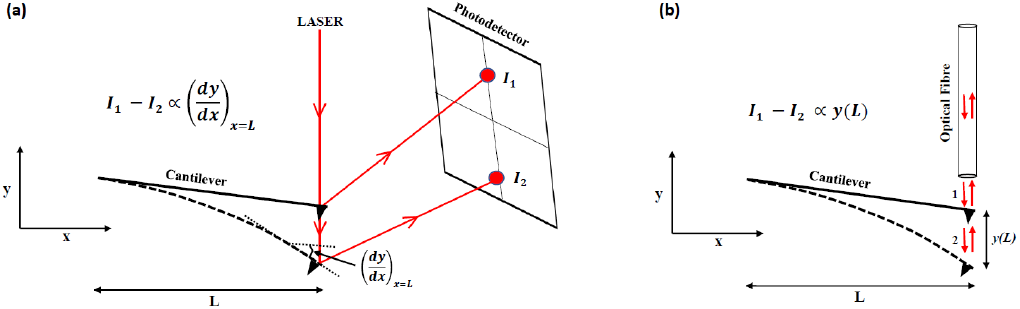
(a) The schematic of the deflection detection set-up. The laser reflecting from the back of the cantilever falls on the position sensitive four quadrant detector. The outputs *I*_1_ and *I*_2_ correspond to the two positions of the laser spot on the detector. This position depends on the bending (*dy/dx*) of the cantilever at *x = L*. The signal from the detector reads the local cantilever bending. (b) In interferometerbased detection, the cantilever acts as one the mirrors in Fabry-Perot etalon. the displacement of the cantilever produces a proportional signal in the photo-diode. The signal from photo-diode measures displacement (*y*) of the cantilever end. In both schematics the cantilever bending and the displacement are exaggerated.

Rectangular cantilevers, made of silicon nitride, from micromasch (Micromasch, Bulgaria) with stiffness 0.05-1 N/m were used for the experiments. Typical dimensions of the cantilevers were length ~ 300 *μ*m, width ~ 30 *μ*m, and thickness ~ 1 *μ*m. Typical resonance frequency of the cantilever in water is ~14 kHz. Cantilever stiffness is calibrated using thermal fluctuation method available with JPK (24). Cantilever base is excited using a dither piezo. A sinusoidal signal from internal oscillator of lock-in amplifier (SRS830, Stanford, California, US) is applied to the piezo and the same signal is used as a reference for the phase sensitive detection. Output signal from the photodetector is supplied to the lock-in amplifier which provides us the amplitude and phase response of the cantilever.

### Protein

Tit in I27_8_ polyprotein constructs containing eight identical domains in tandem were constructed from a plasmid similar to described in reference (25). Expression and purification of the protein was done as described before (26). PBS at pH 7.4 was used as the standard buffer for all experiments. The protein sample (100 *μ*L) in PBS with a concentration of 10 mg/mL was adsorbed onto a freshly evaporated gold coated coverslip assembled in the fluid cell by incubating it on the substrate for 15-30 minutes at room temperature. The sample solution was washed three times with PBS to remove the excess un-adsorbed protein from the working solution. Experiments are performed at low frequencies (~ 100-300 Hz, for off-resonance measurements) and with small amplitudes (~ 1-2 Å). The pulling speed (25 nm/s) kept low compared to the conventional protein pulling experiments. These parameters ensure the linearity of the measurements which is essential to use the existing theoretical models for analysis.

### Theory

In order to describe the dynamics of a rectangular cantilever immersed in a liquid we begin with Euler-Bernoulli beam equation. The equation is solved for sinusoidally excited cantilever. We modelled the hydrodynamics of base-excited cantilever for both in the presence and absence of forces on the tip.

We derive the solution for both bending and displacement of the cantilever at the tip end in off-resonance regime.

The Euler-Bernoulli equation of motion for a homogeneous rectangular cantilever beam immersed in a liquid is

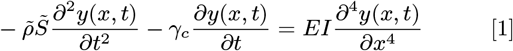

In the model the internal damping of the cantilever has been neglected. The coordinate along the cantilever of length *L* is *x*, where the base is at *x* = 0 and *x = L* is the end/tip-position. Displacement normal to the length at position *x* and time *t* is *y*(*x,t*). The effective mass per unit length is 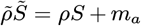, where *ρ* and *S* are cantilever mass density and cross-section area respectively; *m_a_* is hydrodynamic added mass per unit length; *γ_c_* is the cantilever’s hydrodynamic drag coefficient per unit length. *E* is the Young’s modulus of the cantilever and *I* is the second area moment; 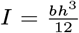 for a homogeneous rectangular cantilever, where *b* and *h* are cantilever’s width and thickness respectively; 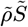 and *γ_c_* depend on the excitation frequency. To solve the equation, we used variable separation method and assumed the solution to be of the form *y*(*x,t*) = *y*(*x*)*e^iωt^*; where *y*(*x*) is the solution of the space part and *ω* is the excitation frequency. Substituting the solution into (1) gives

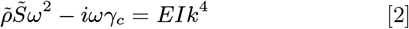

Where the spatial part of the Eq. 1 can be written as

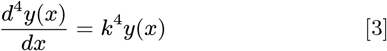

General solution of the above equation is given by

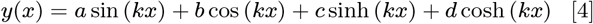

and the slope (cantilever bending) is given by

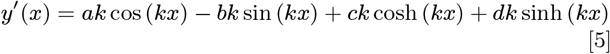

In general,the coefficients *a, b, c*, and *d* are complex numbers and are determined by applying appropriate boundary conditions dictated by the experiments (*y*-displacement, *y*′- slope, *y*′′- internal moment of force, and *y*′′′- internal shear force).

We now define dimensionless parameters, *g* and *z* given by

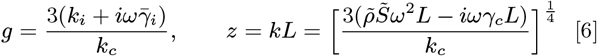

Where *k_i_* and 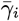 are interaction stiffness and damping respectively; 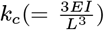 is the cantilever stiffness. These two parameters will be used for implementing approximations for different experimental situations. For low excitation frequencies *z* << 1 and for cantilever having large stiffness *g* << 1.

#### Cantilever excited at base without interaction (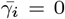 and *k_i_* = 0)

The boundary conditions for cantilever excited from base with amplitude *A* and in absence of any forces acting on the tip are given by

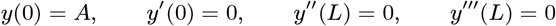

The constants *a,b,c* and *d* in (4) and (5) are determined by using the above boundary conditions. We get the solution for small g and z approximations. (see supporting information theory section for the full derivation)

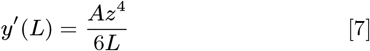

Substituting the value of z that we had defined earlier (see Eq. 6) we get the following solution for the bending of the cantilever at the free end

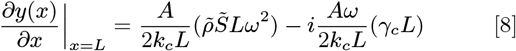

Modulus of (8) gives us the amplitude of the cantilever bending and argument gives the phase difference between drive (or base) and bending at *x = L*

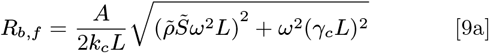

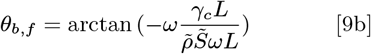

The real and imaginary components are

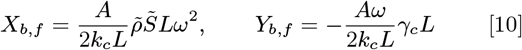

Where subscript *b* stands for bending and *f* for the free cantilever. Constants 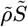 and *γ_c_L* can be determined from free *X* and *Y* signals using (10). Since, *X_b,f_* and *Y_b,f_* are *R_b,f_* sin(*θ*) and *R_b,f_* cos(*θ*) it is important to note that an error-free measurement of *θ*, the phase lag of the tip with respect to base is critical to find the exact values of these constants. The solution for the displacement with *z* << 1

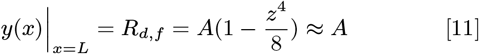

The amplitude of the displacement of the cantilever end (*x = L*) in absence of any interaction is approximately equal to the base amplitude and both move in phase and the phase lag *θ_d,f_* of the cantilever tip with respect to the drive is zero.

#### Cantilever excited at base in presence of interaction (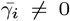 and *k_i_* ≠ 0)

For a cantilever being driven from base and the tip experiencing a linear viscoelastic force due to tethered macromolecule, the boundary conditions are given by

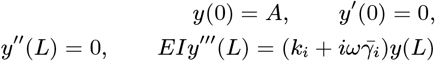

The constants determined using the boundary conditions are substituted into Eq. 4 and 5. Approximations for *z* << 1 and *g* << 1 are then applied to give us (see supporting information theory section for the full derivation)

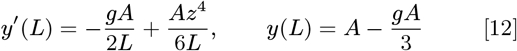

Substituting the values of ‘*g*’ and ‘*z*’ back into the (12) gives the following expressions for slope and displacement respectively

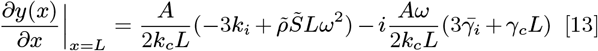

and

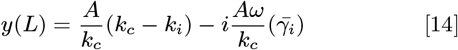

Modulus of (13) gives us the amplitude of the bending (slope) at *x = L*:

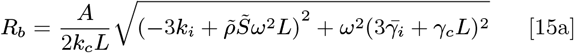

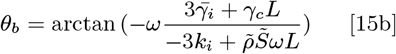

Now the real and imaginary components of the bending are

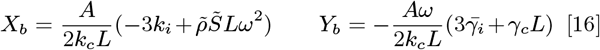

This is identical to the result obtained by Benedetti et al. (7) Further, the amplitude and phase for the displacement *y* in the limit *g* << 1 and *z* << 1 is

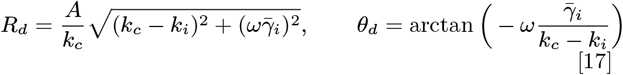

The real and imaginary components of the displacement are

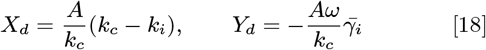

The subscript *d* stands for displacement.

Eq. (16) and (18) are the main results which can be used to estimate stiffness and dissipation. Eq. (16) can be used for experimental data from deflection detection type AFM and equation (18) can be used for interferometer based AFM.

#### Cantilever excited at base and tip in non-deformable contact

Cantilever is excited from the base at *x* = 0 with amplitude *A* and tip is in contact with the substrate. Assuming that both surfaces (tip and substrate) are non-deformable, the boundary conditions are following:

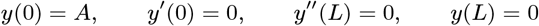

When the cantilever-tip is pressed against a non-deformable surface, the displacement at the tip is zero however bending is nonzero. The slope at x=L:

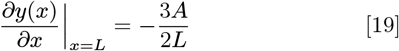

At deep contact region, cantilever bending at *x = L* is proportional to the base displacement and moves 180 degree out of phase. (see supporting information theory section for more details)

### Point Mass model

In an alternative model to describe the cantilever dynamics, the drive force and the force acting on the tip is included in the equation of motion instead of in the boundary conditions.

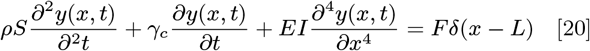

Where *F = F_exc_ + F_i_. F_exc_* is excitation force and *F_i_* is interaction force. Assuming the solution of Eq. 20 *y*(*x,t*) = *y*(*x*)*Y*(*t*). *y*(*x*) could be determined by solving the space part of the Eq. 20 using following boundary conditions:

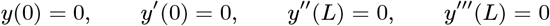

The solution to this equation has all the modes. The point mass approximation considers only the first (or fundamental) mode of vibrations. The cantilever is treated as effective pointmass (*m*\*) attached to a mass-less spring with spring constant *k_c_*) and oscillating at its first eigen-mode resonance frequency. The equation of motion for the oscillating cantilever at its fundamental mode is given by:

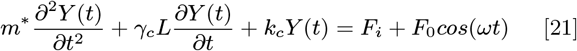

Where 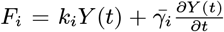 and *F*_0_ = *k_c_A*_0_. *A*_0_ is amplitude when cantilever is far away from the surface and interaction force is absent. The solution of the first mode in presence of the interaction is assumed to represent the entire dynamics of the cantilever adequately (27). The final result which relates the amplitude(*R_d_*), phase(*θ_d_*) to stiffness and dissipation in the off-resonance condition is given by

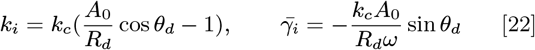

It is to be noted that the solution to the differential equation is for the displacement (*y*) and not the bending (*dy/dx*) of the cantilever. See reference (27) for more details.

## Results and Discussion

### Measurements performed using two detection methods

We performed experiments using both a conventional deflection detection type AFM and a home-built interferometer based AFM. These two detection schemes are not only different ways of measuring cantilever oscillations but more importantly measure different quantities. As described in the methods section, the former one measures bending (*dy/dx*) and the later the displacement (*y*) at the end of the cantilever (see Fig. 1). The amplitude (*R_d_*) of measured displacement(y) and its phase (*θ_d_*) are related to the X-component and Y-component of the simple harmonic motion of the tip by following relations. *X_d_ = R_d_ cos θ_d_* and *Y = R_d_* sin *θ_d_*. Similarly, amplitude (*R_b_*) of bending (*dy/dx*) is related to X and Y component by *X_b_ = R_b_* cos *θ_b_* and *Y_b_ = R_b_* sin *θ_b_*. The X and Y components can be recorded using lock-in amplifier.

Fig. 2 shows the raw data from measurements using a deflection detection type AFM. The I27 octomer whose one end is attached to the tip and other is fixed to the substrate, is pulled away from the substrate at speed of 25 nm/s. The measurements are repeated at two frequencies of 133 Hz and 2 KHz. Fig. 2 (a) and (c) is X and Y components of oscillations in the bending *dy/dx* at 133 Hz and 2 KHz respectively. Fig. 2 (b) and (d) is amplitude and phase of the bending *dy/dx* at the tip end. Before measurements were performed we ensured that i) The phase lag in deep contact is 180 degrees, ii) The cantilever is excited far below resonance so that the phase response is flat and there are no spurious peaks in this region. These two are important criterion for artefact free measurements of dissipation. We observed that for deflection detection type AFM, the cantilever tip oscillates with ~ 6 degree phase lag with respect to the base excitation when immersed in liquid and is freely oscillating. We treat this phase lag as emerging purely from the cantilever hydrodynamics. The Y-component in both these measurements is not varying when the domains are sequentially unfolded. According to Eq. 16, this means that the dissipation is immeasurably low when protein unfolds or the unfolded domains are stretched further. However, the measurement performed at 2 KHz shows that the phase signal shows clear oscillations. Analysis of data in fig. 2 (d) using point-mass model shows that there is measurably high dissipation in single molecule stretching experiments.

**Fig. 2.**
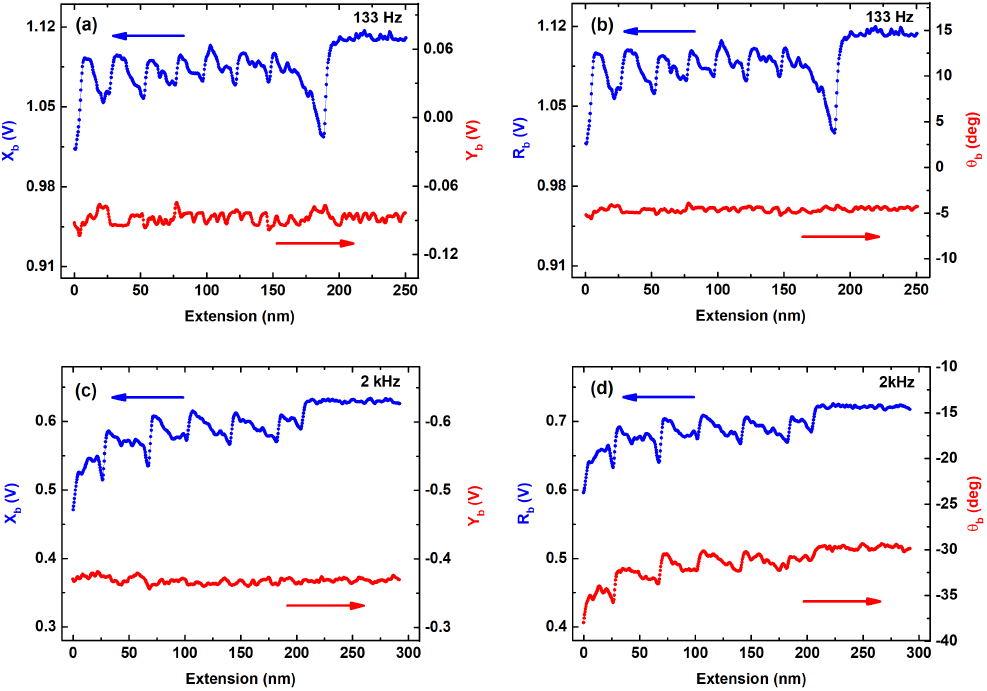
The raw data of the measurements performed using a deflection detection type AFM (a) The X and Y components of oscillations in the bending((*dy/dx*)_*L*_) at the tip end of the cantilever. The measurements are performed at 133Hz. (b)The amplitude and phase of the oscillations in the bending (*dy/dx*) of the tip end of the cantilever. The same measurements are repeated at 2 KHz in (c) and (d). The Y-component in both measurements is featureless and according to Eq. 16 the dissipation in single molecules is immeasurably low. The phase shows clear oscillations in (d), however the corresponding Y-component in (c) does not change as the domains are sequentially unfolded. The use of The point-mass model wherein dissipation depends on phase (Eq. 22) then results in erroneous measurement of dissipation.

**Fig. 3.**
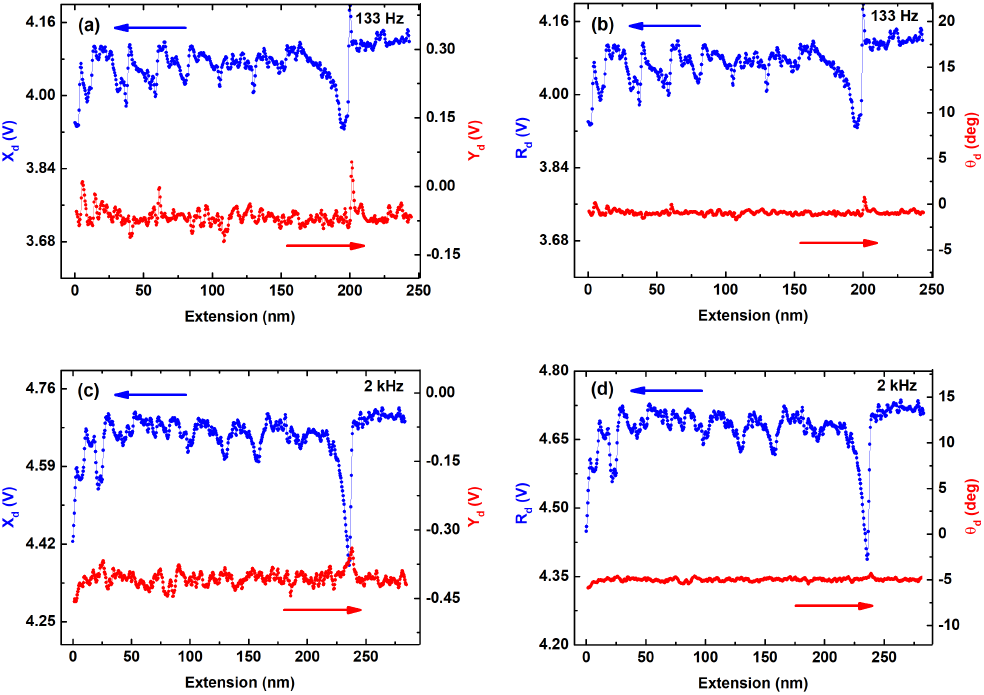
The raw data of the measurements performed using home-built interferometer based AFM (a) The X and Y components of the oscillations in the displacement at the tip end (*y* (*L*)) of the cantilever. Frequency of oscillations is 133 Hz. (b) The amplitude and phase of oscillations in the displacement. (c) and (d). The measurements are repeated for 2 KHz. Unlike the phase of the the bending signal in fig. 2(d), the phase of the displacement does not show variations as the protein unfolds.

Fig. 3 shows raw data of the measurements using homebuilt AFM equipped with interferometer based detection system. This detection measures displacement (y) of the cantilever. Fig. 3 (a) and (b) shows measurements performed at 133 Hz and (c) and (d) shows measurement at 2KHz. We clearly see signatures of protein unfolding under force. The typical measurements shown here at both frequencies of 133 Hz and 2 KHz show that the Y-component of the signal does not show variation while the octamer is unfolded sequentially. Both the amplitude and X-component shows saw-tooth like pattern which is typical of polyprotein unfolding under force. Since the interferometer detection system measures displacement and not the bending in the cantilever, it shows no phase lag when tip is far from the substrate. This is in agreement with result obtained in Eq. 11. Interestingly, unlike the measurement of bending performed with deflection detection AFM (Fig. 2 (d)), the phase of displacement (y) measured using interferometer based AFM does not vary while protein unfolds (Fig. 3 (d)). This means that the use of both continuous beam model Eq. 18 and the point-mass model Eq. 22 yields immeasurably low dissipation when displacement of the cantilever is measured.

Together, Fig. 2 and 3 show measurements of X-component, Y-component, the amplitude and phase of both bending 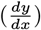 as well as displacement (*y*) signals. Using models used to describe cantilever hydrodynamics Eqs. 16 and 18 in the viscous media these quantities can be used to infer the viscoelastic response of single protein molecules under strain. Clearly, in both measurements the Y-component which is related to the dissipation alone in the molecule is not changing as the unfolded protein domains are pulled with a constant velocity. This indicates that dissipation in the single molecule of titin is immeasurably low. This is in direct contrast with previous claims of measurements performed on single molecules (10, 11, 28).

We estimated the minimum detectable dissipation coefficient from the noise levels in the Y-component signal at our operational parameters (see supporting information section (Minimum detectable dissipation) for more details). It is 5 × 10^−7^kg/s. We conclude that the upper bound on the dissipation coefficient of single unfolded protein is ~ 5 × 10^−7^kg/s. This value is smaller than those reported in the literature.

### Quantification of Stiffness

Further, we used the X-component of the tip-oscillations from both detection methods to calculate the stiffness using Eqs. 16 and 18. It is expected that stiffness determined from these methods should be in agreement with each other. The stiffness should also match with measurements carried out using static deflection of the cantilever in a standard force-extension curve if the dissipation is not present. The derivative of this force-extension curve produces stiffness-extension curves.

To quantify the interaction stiffness from bending measurement data, estimation of the base displacement (*A*), 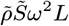, and *γ_c_L* are required. Since the base displacement cannot be directly measured in deflection-detection setup, it can be replaced by the measured bending amplitude at the deep contact using Eq. 19. 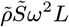, and *γ_c_L* can be determined from the free cantilever *X_b,f_* and *Y_b,f_* signals respectively (see Eq. 10). Finally, for the interaction stiffness and dissipation coefficients, Eq. 16 can be written as:

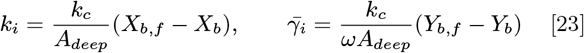

Where *A_deep_* is the bending amplitude in deep contact and *X_b,f_* and *Y_b,f_* is the X-component and Y-component of the the free cantilever without any force on it (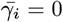 and *k_i_* = 0).

Fig. 4 (a) shows the stiffness-extension calculated from the data in Fig. 2(a) into Eq. 16. The peak stiffness of the unfolded protein is ≈ 20 mN/m. Fig. 4 (b) shows stiffness-extension profile calculated from the data in Fig. 3 (a) and Eq. 18. The stiffness of unfolded protein measured using two detection methods matches well with each other and gives its quantitative estimate for single protein molecule. It is instructive that the stiffness estimated using X-component of the amplitude of bending as well as displacement matches well with static deflection data (see supporting information Fig. S.10). This underlines the robustness of the theoretical modeling used to quantify the stiffness using dynamic AFM having two detection schemes measuring displacement and the bending of the end of the cantilever.

**Fig. 4.**
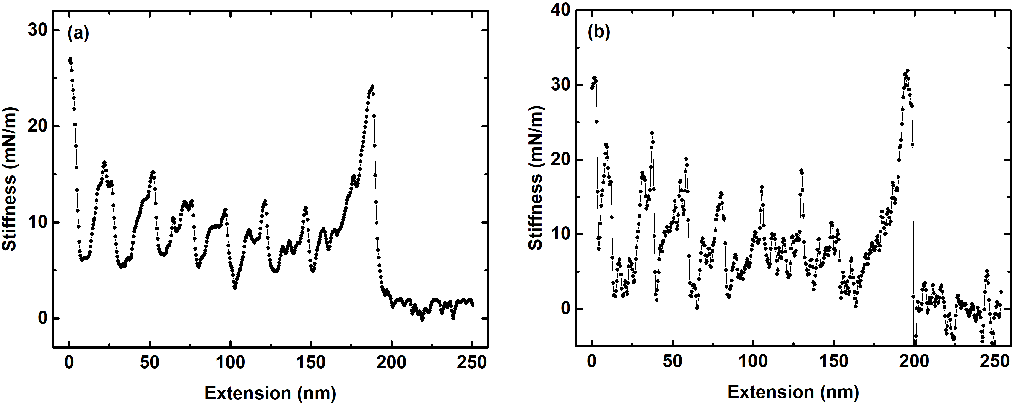
(a) The stiffness estimated from X-component of the bending of the cantilever at the tip end ((*dy/dx*)_*L*_) using a deflection detection measurement. The stiffness is ≈ 20 mN/m. This matches well with reports from other measurement and the derivative of the static force-extension curves (see supporting information Fig. S.10). (b) The stiffness estimated from the X-component of the displacement of the cantilever at the tip end (*y*(*L*)) using a interferometer based AFM. The values of stiffness estimated using both detection schemes are in agreement with each other.

### The effect of extraneous phase

For measurements presented in Fig. 2 and 3, we took care that there is no extraneous contribution to the phase of the untethered free end of the cantilever oscillating in liquid. This is not an easy task for the deflection detection type AFM, since there are variety of intractable sources of extraneous phase. For interferometer based AFM, since cantilever displacement is measured, the phase due to hydrodynamics on the freely oscillating cantilever is zero and it is relatively straightforward to perform measurements with no contributions from such sources. Fig. 5 shows a measurement when such extraneous phase is present. Typically, the phase of the cantilever bending in the deep contact is 180 degrees (see Eq. 19). This is not the case for measurements shown in Fig. 5. The phase in deep contact is ~ 190 degrees. This produces an extraneous phase contribution of the same amount in phase of the freely oscillating lever in the liquid. The Y-component is clearly showing peaks as protein unfolds, and following Eq. 16 one may draw false conclusion about the dissipation in single molecules.

**Fig. 5.**
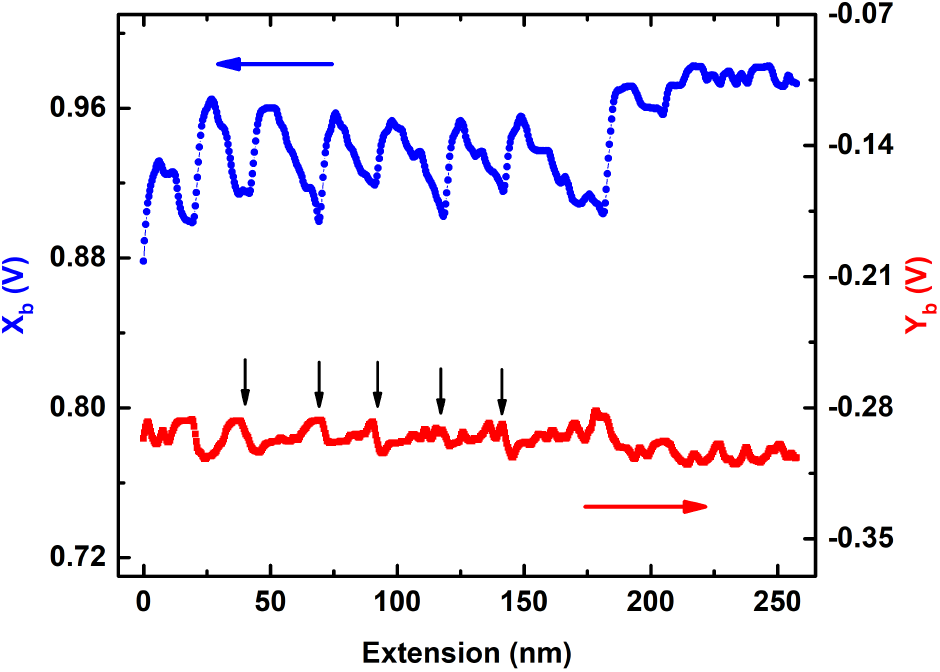
Measurements performed with same cantilever using a different cantilever holder. The phase of the cantilever bending at the tip end is not 180 degree in deep contact (data not shown). The phase of the free cantilever after immersing in the buffer deviates from the measurement in Fig. 2 by the same amount. This is clear case of presence of extraneous phase *θe*. The peaks in Y-signal are identified by arrows. This variation in Y as the protein unfolds is likely to be falsely interpreted as dissipation.

The theoretical models of the cantilever dynamics in the liquid environment, which are presented in the theory section, do not take into account the extraneous phase contributions to oscillating cantilever from variety of sources. In absence of a method to ensure that these contributions are not present, the estimates of stiffness and damping coefficient is not free from errors for the following reason. Let *θ_e_* be the this extraneous phase, *θ_h_* the phase lag due to hydrodynamics and *θ_i_* be phase lag due to molecule’s viscoelasticity (stiffness and dissipation coefficient). *θ_h_* is accounted for in the models. The measured phase when the cantilever is immersed in liquid and is far away from the substrate is *θ = θ_e_* + *θ_h_*. Since *θ_h_* depends on cantilever parameters and viscosity of the medium, it is difficult to estimate the extraneous contribution *θ_e_* to the measured phase lag and compensate for it in the experiments.

In the following we explain how the presence of extraneous phase leads to changes in Y-component of the amplitude of bending (*dy/dx*) due to variation in molecule’s stiffness alone while it is being stretched.

We have *θ = θ_i_* + *θ_h_* + *θ_e_*.

By substituting the phase lag due to protein’s viscoelastic response and the cantilever hydrodynamics from Eq. 15, the total phase lag *θ* is given by

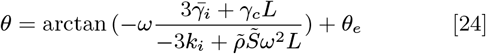

The first term is the phase lag in the cantilever deflection due to both the molecular stiffness and the dissipation as well as the hydrodynamic damping. The extraneous phase can not be included in the model and it sits outside the X and Y components due to hydrodynamics and molecular viscoelasticity. Let’s assume that molecule being stretched is purely elastic entity with dissipation 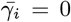. Since the molecular stiffness is entropic, it changes with extension. Fig. 6 (a) graphically represents Eq. 15 and 16 in the absence of extraneous phase. The amplitude and phase of the cantilever bending changes due to variation in molecule’s stiffness. The phase lag will have contribution *θ_i_* from the molecule’s stiffness and the amplitude changes from *A* to *A*′. Since the dissipation is zero, the Y-component (*ωγ_c_L*) does not change. This should only affect the X-component and it reduces from 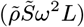 to 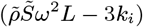.

**Fig. 6.**
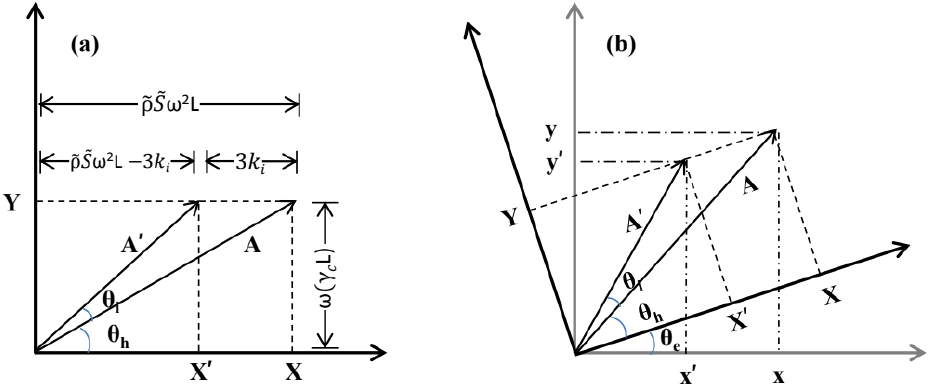
Schematics to show how extraneous phase produces artefacts in the measurement. (a) *θ_e_* = 0, 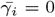; The change in *k_i_* affects both phase and amplitude but Y-component remains unchanged. (b) When *θ_e_* is nonzero, The phase and amplitude changes but Y remains unchanged in the co-ordinate system XY wherein *θ_e_* = 0. However in the xy coordinate system, the change in stiffness of the molecule *k_i_* changes the Y-component from *y* to *y*′. Using Eq. 16 this can be misinterpreted as the dissipation. The change in Y-component is due to change in molecule’s stiffness.

Consider now a situation wherein there is extraneous phase contribution (*θ_e_*) to phase when the cantilever is far from substrate and is freely oscillating. Fig. 6 (b) graphically shows the situation when molecule’s dissipation is zero and there is extraneous phase *θ_e_*. The xy is the coordinate system wherein the total phase lag is *θ = θ_i_ + θ_h_ + θ_e_*. The coordinate system XY is rotation of coordinate system xy by *θ_e_*. In this co-ordinate system the phase and amplitude changes purely due to the molecule’s stiffness in such a way that Y-component does not change. However, in xy coordinate system there is change in Y as shown in the figure due to this additional phase lag *θ_e_*. This change in Y may get wrongly interpreted as the dissipation signal. This arises from having a extraneous phase *θ_e_* which is not considered in the model. It is difficult to theoretically predict the hydrodynamic phase lag of the cantilever. It depends on the cantilever stiffness and viscosity of medium and hence is not known *a priori*. It can not be compensated in order to have phase lag emerging from hydrodynamics alone. In this context, the interferometer based experiments are crucial. The theory predicts that the cantilever displacement has zero phase lag with respect to the excitation signal due to hydrodynamics. This is also verified by experiments. Any additional phase lag due to spurious peaks or other sources can be easily identified and care can be taken to avoid cross-talk between the stiffness and dissipation channel.

The above discussion suggests that if there is significant phase lag due to hydrodynamics alone, the phase may have oscillations as the protein unfolds since this large hydrodynamic phase lag makes the numerator inside the arctan bracket large. In such situations the theory predicts that features may appear in phase data even if the dissipation coefficient 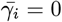. This is because the stiffness is non-zero and is varying under strain. We performed experiments by keeping all other parameters of the experiment same, except viscosity of the medium. Fig. 7 shows a protein pulling experiment after addition of glycerol in the buffer which enhances viscosity. This makes the term *γ_c_L* large and thereby the numerator inside the arctan bracket is also large. In this situation, the *θ_h_* is significant (~ 18 degrees). It was ~ 6 degrees for the same cantilever and much less viscous buffer. As a result, both the phase and Y-components do not change as the protein unfolds. However, after addition of glycerol, the change in molecule’s stiffness when it is stretched affects the phase as molecule sequentially unfolds, but the Y-component does not change from its value which was for freely oscillating lever.

**Fig. 7.**
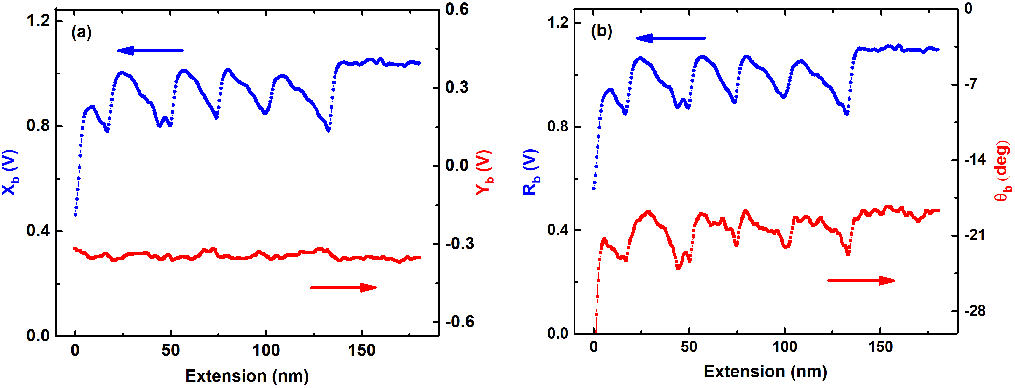
Measurements performed with 50 *%* glycerol in the buffer. This increases viscosity of the medium by ~ 8 times. The *θ_h_* is nearly 18 degrees, however *θ_e_* = 0. As the polyprotein is pulled and the domains unfold, the phase changes but the Y-component does not show any variation. This shows that even if the phase is far from zero due to hydrodynamics alone, the Y-component shows no variation indicating again that the dissipation coefficient is immeasurably low. This is in agreement with results in Fig. 2 (b) and (d).

In a separate set of experiments, we added external phase to the cantilever oscillating in liquid and performed the unfolding experiments. It is seen that Y-component starts to show features corresponding to unfolding events. We performed measurements wherein there is possibility of extraneous phase contributions from electronic or other sources and the Y-component is not free from features (see supporting information section, Fig. S.11).

These experiments clearly show that even if there is a large phase lag due to hydrodynamics when the cantilever is freely oscillating in the liquid, it does not introduce artefacts while interpreting Y-component as a dissipation signal. This is because the models take into account such phase lag due to cantilever hydrodynamics. However, if there is extraneous phase added to the measurement which is not accounted for in the models, one certainly sees variation in the Y-component which is not due to dissipation of the unfolding protein.

Benedetti et al. in their work have warned against the use of point-mass model and interpreting phase signal as dissipation in single molecules. They have shown that the Y-component of the oscillations in deflection (*dy/dx*) as the signal which contains the information about the dissipative processes. We stress here that even the Y-component is not free from errors as suggested by them if the extraneous phase is not completely removed from the measurements. The sources of this phase are listed in supporting information section.

### off-resonance operation

Besides *θ_e_*, an experimenter has to safeguard against other sources of artefact. The most important is choice of operational frequency in the off-resonance regime. We ensured that our measurements were *truly* off-resonance. The parameter *z* defined in theory section determines off-resonance operation. As shown in theory section, at large frequencies, the higher order terms in the solution for *y*′ need to be considered. It makes the solution for amplitude(*R_b_*) and phase (*θ_b_*) more complicated than Eq. 15. In the limit of *z* << 1 and for a given cantilever and viscosity, the X and Y components entirely depend on stiffness and dissipation respectively as seen in Eq. 16. For large frequencies, however the X and Y component both have contributions from stiffness as well as dissipation coefficient of the molecule. In order to calculate stiffness and dissipation from experimental data at higher frequencies where z ~ 1, Eq. 16 can not be used. We performed measurements at higher frequencies and we observed clear peaks in the Y-component data (see supporting information, Fig. S.15). They appear due to changes in the stiffness of the molecule. It is of paramount importance that measurements are performed at truly off-resonance condition if one wants to use X-component alone to determine the stiffness and the Y-component to determine the dissipation coefficient. (see supporting information section, Eq. S.35)

### Point-mass model and displacement detection

We have presented data from both, a more conventional deflection detection as well as interferometer based detection schemes. The cantilever equation of motion has different solutions for bending ((*dy/dx*)_*L*_) and displacement(*y*(*L*)) given by Eqs. 16 and 18. The displacement *y* does not have any phase lag when immersed in liquid whereas *dy/dx* has hydrodynamic phase lag due to viscous medium (see Eqs. 9 and 11). We have already pointed out that measurement of displacement has an enormous advantage over measurement of bending (*dy/dx*) because the extraneous phase lag is easily identified to avoid artefacts. It has another important advantage. We show in this section that a simple minded point-mass model, which suggests that phase and amplitudes are related to the dissipation and stiffness is successful in quantification of viscoelasticity at nano-scale in liquid environment. Many important claims in the past have used point-mass model to quantify stiffness and dissipation of molecular layers from the amplitude and phase (1, 3, 4, 29). The discussion in this section shows that dissipation estimates of liquid layering using point-mass model are accurate.

The solution to Euler-Bernoulli equation of motion for a homogeneous rectangular cantilever beam immersed in a liquid is given by Eq. 14 and the X and Y component of this displacement oscillations is given by Eq. 18. The pointmass model assumes the cantilever to be a point-mass with a mass-less spring. The stiffness and dissipation is related to the change in amplitude and phase by Eq. 22. According to raw data in Fig. 2, the use of point-mass model would suggest that the molecule being stretched under external force has measurably high dissipation coefficient. This is the common error made in many claims of viscoelasticity of single macromolecules (10, 11). The raw data of the interferometer based measurements shown in Fig. 3 reveal that both the phase and Y-component are featureless when polyprotien unfolds sequentially, whereas the phase has features in the deflection detection data in Fig. 2 (d). Further, we used both the pointmass model Eq. 22 and results from continuous rectangular beam Eq. 16 to the data in Fig. 3. We quantified stiffness using the amplitude and phase data (Fig.3(b)) in Eq. 22 and the X-component data (Fig. 3 (a)) in Eq. 16 and compared the two. The inset shows part of the data wherein the unfolding is clearly seen. Interestingly, both models give the exact same values of stiffness as seen in the stiffness extension curves in the inset. The blue points are stiffness values calculated using continuous rectangular beam model and red points calculated using point-mass model. For large values of stiffness wherein the amplitude changes are high, the two start to deviate from each other. The point of deviation is 10 percent change in amplitude. It shows that if the change in amplitude is within 10 percent, one can safely apply the point mass model Eq. 22 to the amplitude and phase data and can accurately estimate the stiffness of nano-scale entities as well as single molecules. Supporting information Fig. S.13 shows plot of both Eqs. 18 and 22. It is seen that the two solutions start to differ from each other after 10 percent change in the amplitude.

Since the protein unfolding experiments did not show any detectable dissipation, we performed similar measurements on water confined to nanoscale. Such nano-confined water shows layering of water molecules parallel to the confining walls. Previous measurements using our interferometer based AFM have revealed that nano-confined water undergoes a dynamic solidification (4). The measurement of dissipation coefficient had been crucial for drawing conclusions about the nature of such molecular layers of water. Here, we repeated the measurements on water layers using interferometer-based AFM. The water is confined between two parallel surfaces, one surface being the cantilever-tip and other freshly cleaved mica. Before the experiment, cantilever-tip is scanned on mica surface to make it flat and smooth. When the confinement is below 1 nm, the behaviour is different compared to the bulk water due to layering. We observed an oscillatory behaviour in X-signal (stiffness) as well as in Y-signal (dissipation coefficient). Fig. 9 shows the peaks in dissipation coefficient separated by molecular diameter of water. The Average separation between the peaks is ~ 2.52 Å which indicates formation of layers in the confinement region. Experiment is done with 8 Å/s approach speed and the observed dissipation in two to three layers of water is ~ 10^−4^*kg/s*, which is consistent with the previous report (4). It is important to note that the phase of the displacement was zero degrees as theoretically expected for the cantilever immersed in liquid and far from the substrate. It peaks at separation between tip and sample which is integral multiples of molecular diameter (see supporting information Fig. S.18). The variation in phase signal is not observed in case of the protein unfolding experiments. Comparing the results of experiments performed on protein and water using the same instrument suggests that the dissipation can be measured and accurately quantified using the proposed method if it is above the detection limit of the AFM. Such a limit can be obtained by knowing the detection sensitivity of AFM and the thermal noise of the cantilever. The measurements confirm that the dissipation in the single unfolded protein molecule is below the detection limit and hence could not be observed. Further, like the quantification of stiffness using point-mass and the continuous rectangular beam models, we compare the two models used to quantify dissipation coefficient from the measured data. Fig. 9 shows dissipation coefficient obtained using both models match with each other exactly. Together, Fig. 8 and 9 clearly show that one can use the point-mass model to accurately calculate stiffness and dissipation coefficient of nanoscale systems from the measured data provided the displacement of the cantilever is measured. We have calculated dissipation coefficient of water layers using point-mass model following the procedure described in (1).

**Fig. 8.**
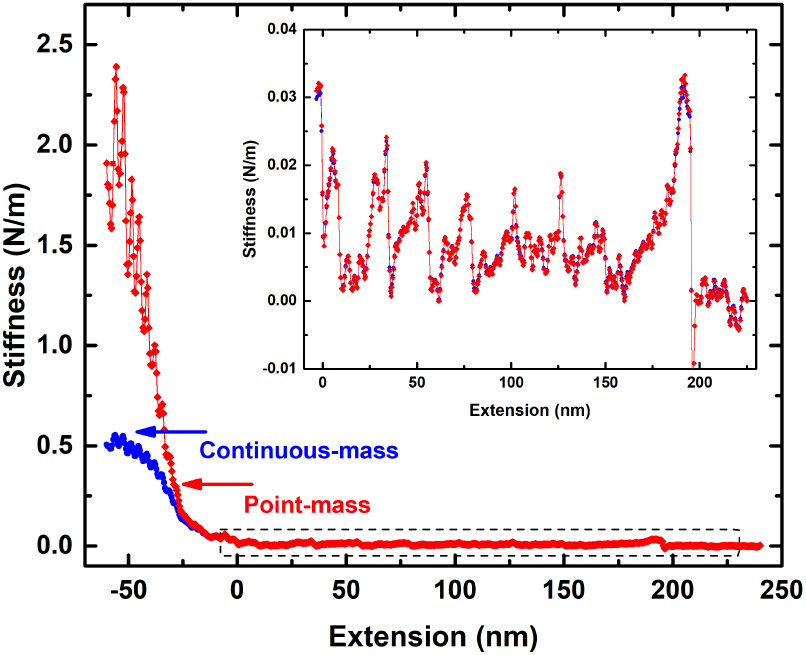
Stiffness calculated using the X-components of the displacement (data in Fig. 3 (a)). The blue curve is stiffness-extension using continuous beam model Eq. 18 and red curve is stiffness extension using point-mass model, Eq. 22. For large stiffness values where the amplitude of the displacement changes significantly, the two curves deviate from each other. The inset shows the magnified data from the region shown by the dotted rectangle wherein polyprotein unfolding is clearly seen. Both the models yield same results in this region where amplitude does not change significantly.

**Fig. 9.**
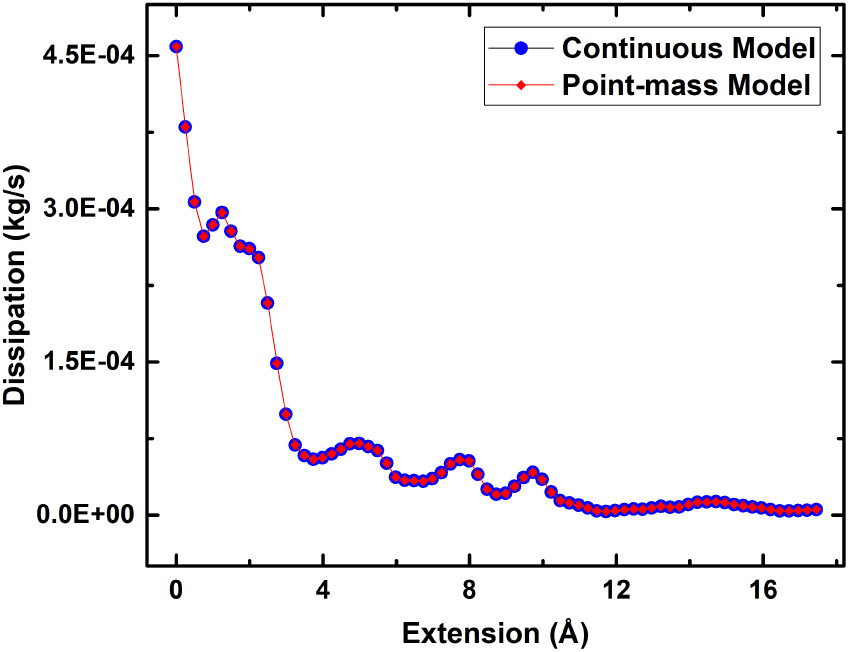
The dissipation coefficient computed using both models given by Eqs. 18 (red filled circles) and 22 (blue filled circles) and the phase and Y-component of the displacement amplitude shown in supporting information Fig. S.18. Dissipation shows clear oscillations peaked at separations equal to diameter of water molecules. Both approaches of analysing the data give identical results. The variation in phase and Y-component is not seen in case of proteins.

### Dissipation in single macromolecules

In literature there are numerous claims of measurement of viscoelasticity of single molecules using AFM (9–11, 28, 30–33). The early reports have shown that the viscoelasticity is dominated by elastic response and the dissipation could not be detected for nucleic acids and Poly Ethylene Glycol (34, 35). However, there are many other reports of measurement of dissipation coefficient for single molecules of protein, flexible polymers and polysaccharides (10, 11). A closer look at the experimental details reveal that these measurements were performed at higher frequencies, sometimes very close to resonance, and with much larger amplitudes (2 −10 nm). They are not truly off-resonance. In our opinion, the simple approximation of the cantilever dynamics to a point-mass model is not adequate to interpret the experimental data performed at higher frequencies or close to resonance. We have clearly shown in our experiments that the measurements are prone to artefacts if phase data is used either at same viscosity and higher frequencies (see supporting information Fig. S.15), or low frequencies and higher viscosity (see supporting information Fig. S.16). Both these situations clearly show features in the dissipation data due to cross-talk between stiffness and dissipation coefficient of the molecule. There is no mention in these experiments if the phase is offset by a certain value while performing experiments. If this offset is not carefully provided then the phase data produces artefact in determining the dissipation coefficient. Moreover, if the phase and amplitude of cantilever *bending* is measured, then the point mass approximation can not be used to analyse the data. This point has been recognized earlier by many (7, 16, 17, 36–41). We show here that the point-mass approximation is useful if one is measuring the displacement of the cantilever with interferometer-based detection scheme.

Higgins et al. used frequency modulation AFM to directly measure stiffness of I27_8_. Unfortunately the dissipation is not measured in these experiments.

Khatri et al. measured the viscoelasticity of titin I27_8_ homopolymer by measuring the thermal fluctuation of the cantilever wherein the tip is tethered with the molecule and held at a fixed force (28). They observed stiffness (≈ 40-80 mN/m) and dissipation coefficient (≈ 2 × 10^−6^ kg/s) of the unfolded chain. This is a different type of measurement. It is important to recognize that point-mass approximation is used to model the cantilever dynamics and the thermal fluctuation of cantilever bending is measured. Our measurements advise-caution while interpreting the constant velocity pulling of such approximation. The analysis of thermal fluctuation data by considering the full details of cantilever’s geometry is complicated and future investigations may reveal if this method is actually measuring dissipation in single molecules.

Using both types of detection schemes to measure cantilever response we concluded that the dissipation coefficient is immeasurably low. The estimate of minimum detectable dissipation coefficient and stiffness using both methods is useful in determining the upper limit on the value of dissipation coefficient for the unfolded chain of I27_8_. This in turn allows for determining a upper bound on Maxwell’s relaxation time on the unfolded chain.

In conclusion, we performed measurements of viscoelasticity of single protein (I27_8_) using both the deflection detection type AFM and interferometer based AFM. We have observed that the dissipation in macromolecule is immeasurably low for AFM’s detection limit. However, with interferometer based AFM and similar experimental parameters, it was possible to measure dissipation in molecular layering of water under nanoconfinement. The estimates of stiffness of macromolecule, using both detection methods is in agreement with each other and the static force-extension measurements. The results imply that using phase signal of cantilever bending along with point-mass approximation of the cantilever hydrodynamics leads to artefacts in the dissipation measurement of single molecules. We demonstrated that with appropriate care, the simple point-mass model used for measurement of viscoelastic response at the nanoscale in liquid environments is adequate if displacement, and not the bending, of the cantilever is measured. It means that the phase of the cantilever displacement can be interpreted as the dissipation signal. The results have a strong bearing on the time-scales of initial collapse in folding of I27 domains and is an important step towards understanding the passive elasticity of titin in muscles.

## Supporting Information Appendix (SI)

### Theory

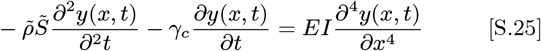

The spatial part of Eq. S.25 equation can be written as

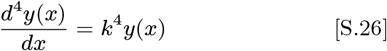

General solution of Eq. S.26 is:

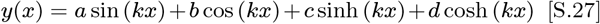

and slope

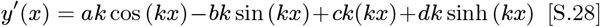

### Base excited and tip-sample interaction is present

Boundary conditions:

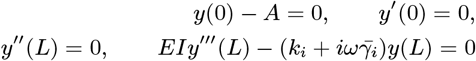

Above set of linear equations can be written in matrix form: *MX − N* = 0 and solve for *X* using matrix method.

Where M and N are

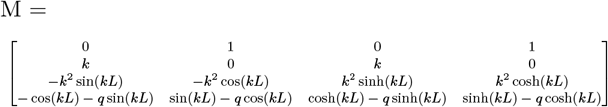

and

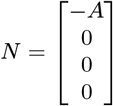

Where 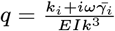.

Solving the matrix equation *MX − N* = 0, the coefficients (*orX*) are following:

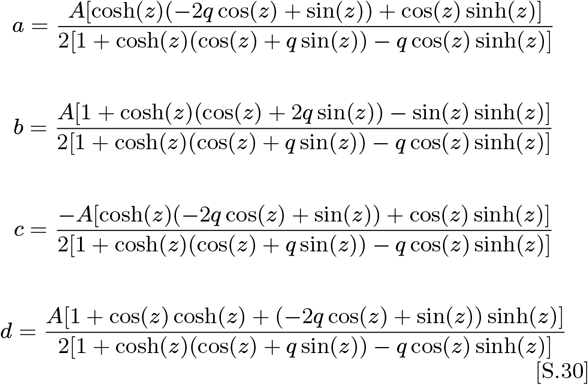

Where *z = kL*.

Solution (displacement) at *x = L* can be written as:

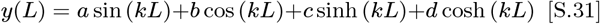

and slope

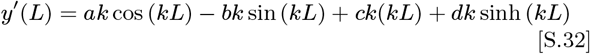

Substitute values of *a, b, c*, and *d* into Eq. S.31 and S.32. Take the Taylor expansion for small *z*^4^ and *g*, displacement and slope are following:

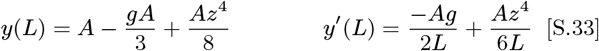

Whereas, full solution with higher order terms are as following:

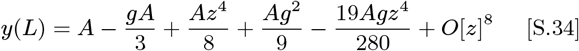

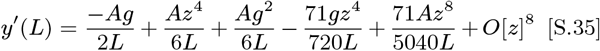

Consider first two higher order terms in Eq. S.35:

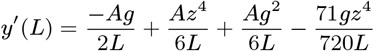

Real and imaginary parts are:

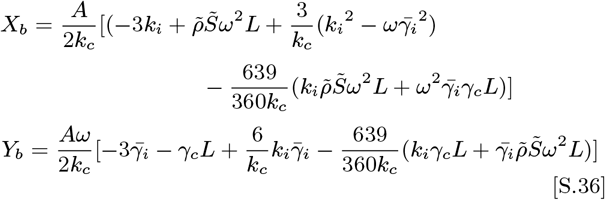

In Eq. S.36 Y contains interaction stiffness which could lead variation in the Y-signal due to variation in the interaction stiffness even in absence of dissipation.

Applying the appropriate boundary conditions, similar procedure could be followed to derive the solutions for the freely oscillating cantilever and cantilever is in non-deformable contact with the substrate.

**Fig. S.10.**
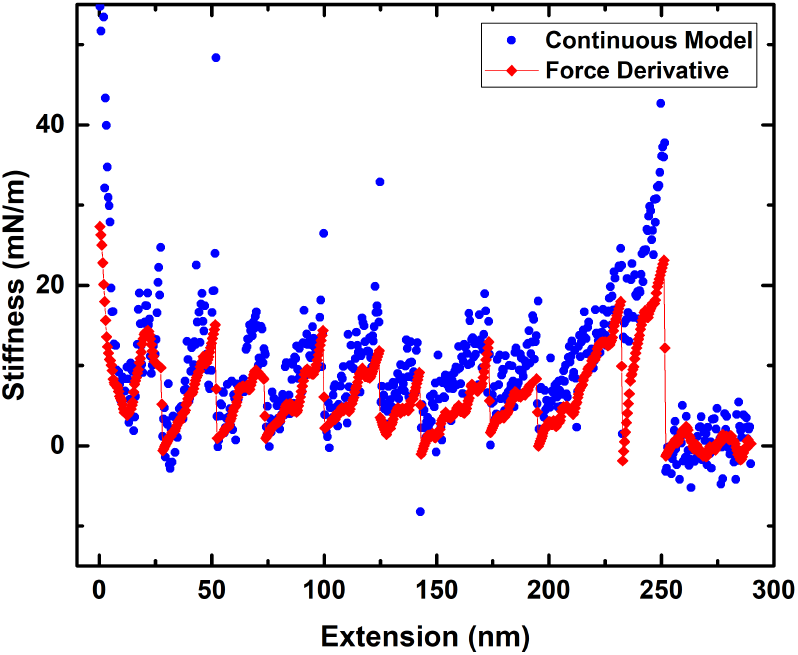
Directly measured stiffness using deflection detection scheme and the derivative for force simultaneously measured by static deflection of the cantilever

### Quantification of Stiffness

Fig. S.10 shows directly measured stiffness using deflection detection scheme and the derivative for force simultaneously measured by static deflection of the cantilever. The two measurements give identical result. This also indirectly means that the dissipation coefficient is zero.

### The effect of extraneous phase

Fig. S.11 shows the measurement performed by adding phase to the cantilever. Without adding the external phase the cantilever end shows a phase lag of *θ_h_* ~ 6 degrees. This is due to the hydrodynamics and the Y-component is free of any variations as as shown in Fig. 2 (b). Any addition to this phase externally using phase shifter produces variation in Y-signal as the molecule is stretched and domains unfold one after the other. It is also seen that the as added phase is close to 90 degrees, the X-signal shows no variation whereas Y shows maximum variation when molecule is stretched. This indicates that for any uncompensated nonzero *θ_e_*, the Y-signal shows a saw-tooth like pattern when the octamer is pulled on and unfolds sequentially. Thus, if the a extraneous phase is not correctly compensated for or completely removed from the measurement, one can wrongly interpret the Y-signal as a measurement of dissipation in single molecules when they are stretched.

### sources of extraneous phase *θ_e_*

Following are the sources of extraneous phase when the measurements are performed wherein the base of the cantilever is excited.

1. Electronic phase: The cantilever is excited by providing a electrical signal to a piezo excitor on which the base of the cantilever is mounted. The deflection (*dy/dx*) or the displacement (*y*) of the cantilever’s tip end is measured by optical means. A current to voltage converter is then used to amplify the signal to make it measurably large. Typically a high gain-bandwidth amplifier is used for this purpose and at reasonably low excitation frequencies (0.1 - 1 kHz), there is no delay between input and output of this amplifier. The main source of extraneous phase is then from the electrical connections made to the piezo used for excitation of the base. It is straightforward to measure this phase contribution by recording the phase lag in deep contact. Any deviation from 180 degrees can be attributed to electronic phase contribution. Furthermore, the absence of electronic phase offset can be confirmed by observing the phase lag in air. It is well known that the cantilever tip has no phase lag when oscillating in air in the off resonance regime (see (27)). We observe zero phase lag in air and 6 degrees phase lag in water. Since in air there is no phase contribution due to electronics, the cantilever immersed in liquid is also devoid of any extraneous phase due to electronics or electrode connections to the piezo-excitor.
2. Local peaks far from the cantilever resonance: Fluid borne excitation: Xu and Raman have investigated different methods of cantilever excitation and have pointed out that cantilever base excitation using a piezo results in tip oscillation having two contributions. The one is structure-borne excitation and the other is fluid-borne excitation. The fluid-borne excitation are due to exciting the local fluid surrounding the cantilever and the base which is roughly thousand times bigger then the cantilever itself. These excitation are due to oscillations of the chip on which the cantilever is mounted and strongly depend on the geometry of the liquid cell. This typically results in “forests of peaks” in cantilever excited in liquids. We performed measurements when the frequency sweep is not free from such peaks. Fig. S.12 shows result of such a measurement. The arrows point at the peaks in Y-signal which match with unfolding of protein where the stiffness peaks. These peaks are due to variation in the stiffness as shown in the schematic of Fig. 6 (b) and not due to dissipation in the molecule.

**Fig. S.11.**
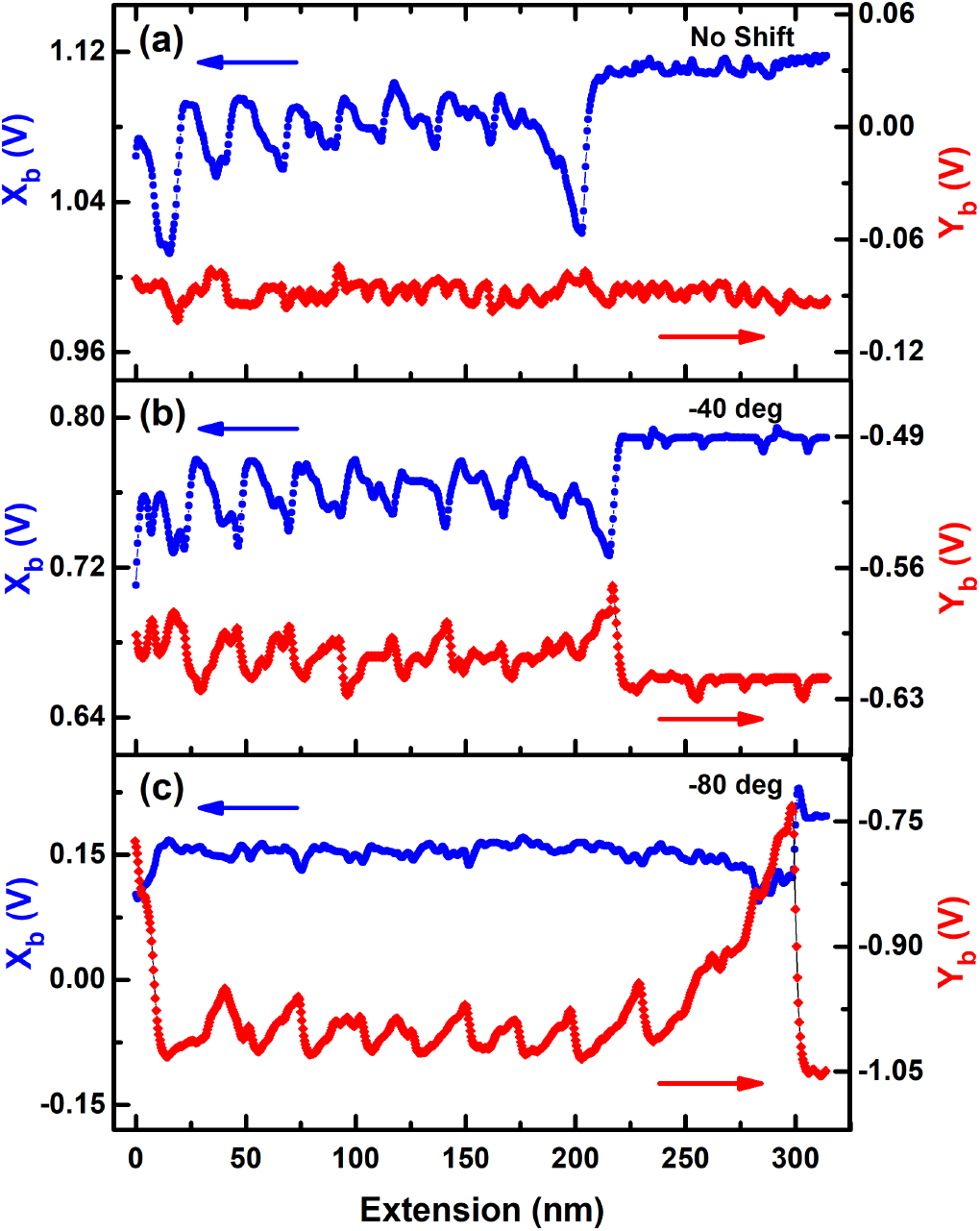
Three unfolding experiments carried out with (a) no added electronic phase (b) by addition −40 degree phase (c) addition of −80 degree. There is no variation in Y without addition of external phase. The variation is seen in Y with addition of external phase and close to −90 degree the there is more variation in Y-component as compared to X-component.

**Fig. S.12.**
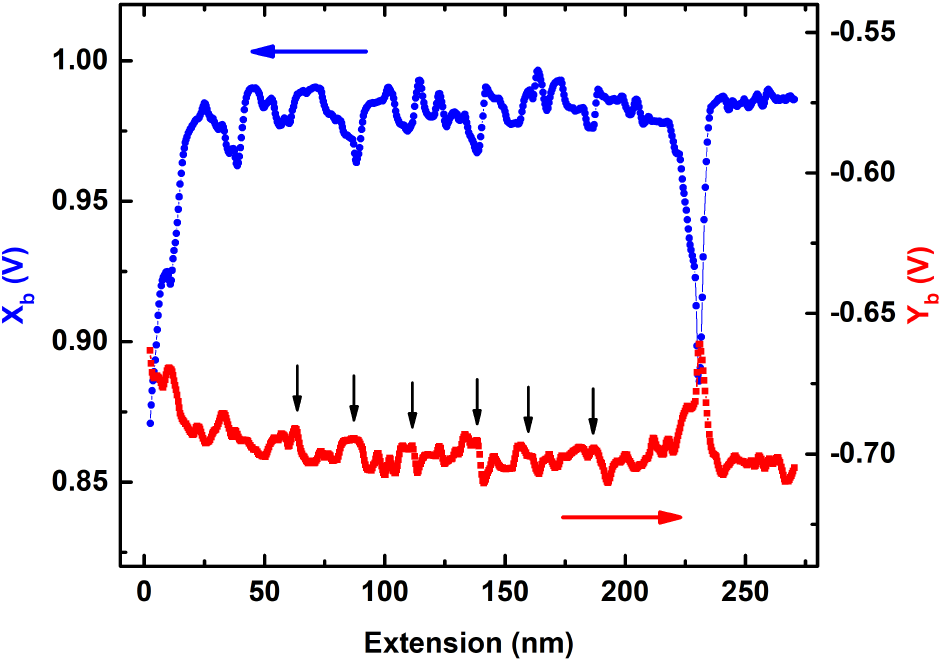
Measurements performed when the frequency sweep is not free from local peaks. The Y-signal shows variation and peaks where the stiffness is maximum. This may wrongly get interpreted as the dissipation signal.

It should be ensured that for *true* off-resonance there are no local spurious peaks. This adds phase to the end of the cantilever and contributes to *θ_e_* Fig S.12 shows a pulling experiment wherein there is a local peak near the excitation frequency ~ 100 Hz. This spurious peak produces a phase lag that is not accounted for in the modeling and thus contributes to *θ_e_*. It can be clearly seen that the Y-signal now shows variation while octomer is sequentially unfolded. This is an artefact due to extraneous phase.

### Off resonance operation

In the following we discuss off-resonance operation and use of Eqs. 16 and 18 to estimate the viscoelastiy of single molecules from the data.

The quality factor of cantilever resonance in air is typically around 100-1000 and in the vacuum environments it is 10,000-100,000. The off-resonance operation is ensured in these situations in a relatively straight-forward manner and nanoscale visco-elastic measurements are free from errors. However, the quality factor in liquids is of the order of 1-5. This poses a serious problem in ensuring that the cantilever is excited at a *truely* off-resonance frequency. Typically, the cantilever can be excited around 100 Hz and can be treated truely off-resonance, if first resonance of the cantilever is around 10 kHz.

**Fig. S.13.**
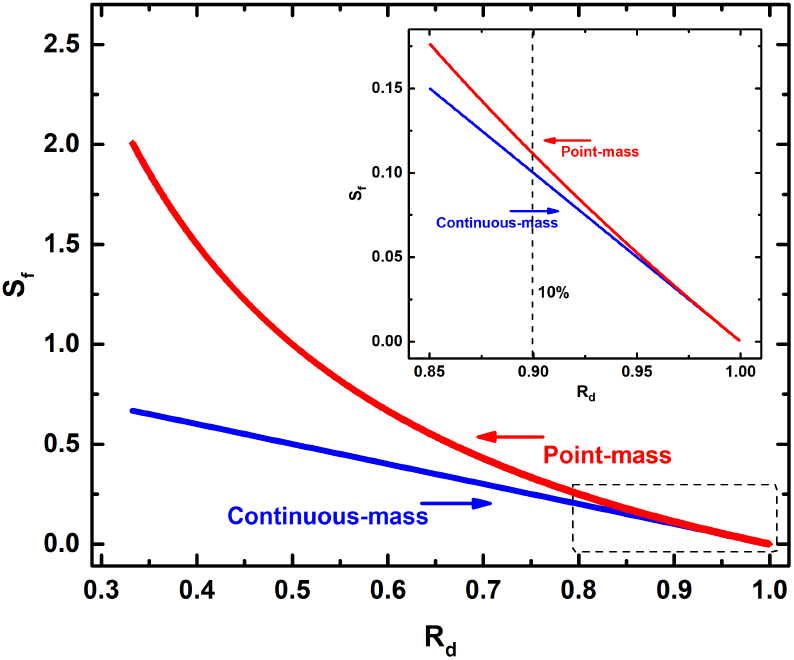
The solutions of point-mass model and homogeneous rectangular beam are plotted together. It is seen that when the amplitude change due to tip-sample interaction or the viscoelasticity of the macromolecule is less than 10 percent, the values of stiffness given by Eq. 18 and 22 are identical. Above 10 percent change in amplitude, the point mass model overestimates stiffness values.

**Fig. S.14.**
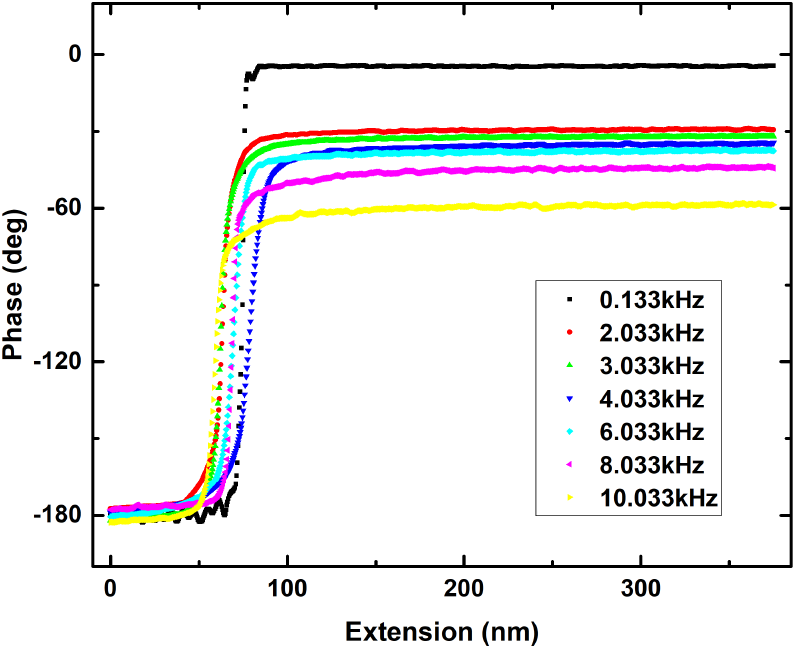
The phase data at different frequencies. The curves are taken when the cantilever engages with the substrate and the free cantilever moves into deep contact. The phase of the free cantilever varies with frequency. In deep contact the phase lag is always 180 degrees independent of drive frequency.

We performed experiments to test the range of off-resonance frequencies where one can safely operate and use Eqs. 16 and 18 to quantify viscoelasticity from the experimental data and measurements remain free from artefacts. Using the same cantilever which were used to perform experiments of Fig. 2 and 3, we operated at higher frequencies to record X and Y components of the cantilever oscillations in both deflection detection AFM and interferometer based AFM. At relatively high frequencies, the variation is observed not only in phase but also in Y-signal. Fig. S.15 shows Y-component recorded using both detection schemes. At 8 KHz, we clearly see variation in the Y-signal as the octamer unfolds.

**Fig. S.15.**
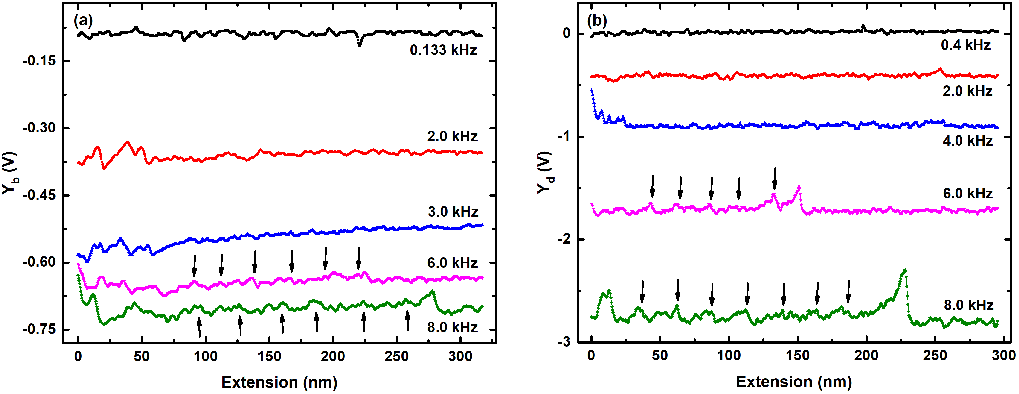
(a) Measurements performed at higher frequencies using deflection detection type AFM. The Y-component of the bending oscillations shows variations after 4 KHz. (b) Measurements at higher frequencies using interferometer based AFM. The Y-component of the displacement oscillations shows peaks at extensions where domains unfold. In both measurements, the phase response is not flat. For higher frequencies the phase and the Y-component is not constant when the tip is far from the substrate. In these measurements the electronic phase lag was ensured to be absent by recording phase in the deep contact region (see Fig. S.14).

Secondly, we changed the viscosity of the solvent in which the experiments are preformed by adding Glycerol (70%) to it. This changes the viscosity of the medium by more than 30 times. We used similar operational parameters used in Fig. 2 and 3. The experiments were performed with similar parameters as was used in low frequency measurement case (frequency = 133 Hz, speed = 25 nm/s, cantilever stiffness = 0.73 N/m). We observed the Y-signal and phase signal when the single molecule is stretched (see Fig. S.16). This could be misinterpreted as the dissipation from the stretching molecule. In this highly viscous medium the cantilever quality factor goes down to much less than ~ 1 and the system is overdamped. The drive frequency (133 Hz) is not really off-resonance in this case. The frequency sweep clearly shows no resonance peak and around 133 KHz, the cantilever response is not flat (see Fig. S.17).

**Fig. S.16.**
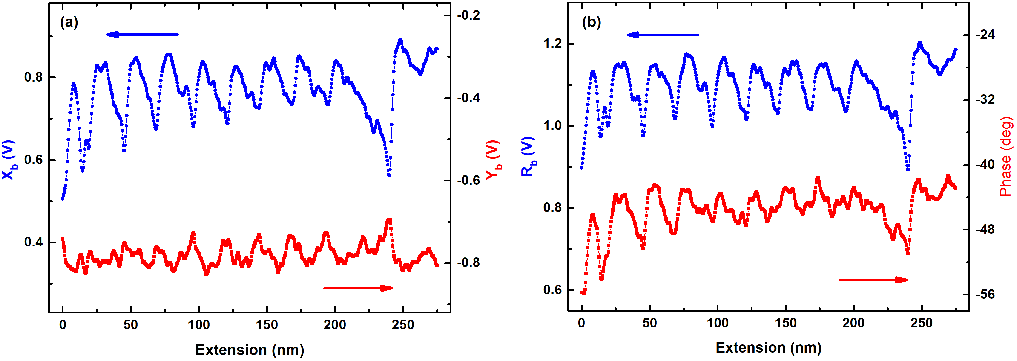
Measurements performed in 70% glycerol in the solution using deflection detection scheme. At higher viscosity, the measurement at ~ 100 Hz is not truly off-resonance. The Y as well as phase shows variation as protein is unfolded sequentially.

In both these measurements, the observed variation of Y-signal, which represents the dissipation alone (Eq. 16), is not due to the dissipation but it is result of the crosstalk between stiffness and dissipation. It is seen from Eq. S.36 that molecular stiffness appears in Y if we consider the higher order terms were considered. This clearly shows that at high frequencies (1 KHz) but still away from resonance (14 Khz) the X and Y components of the tip oscillations both contain stiffness and dissipation and produces a cross-talk in the measurement. It is of paramount importance that the operation is *truely* off-resonance.

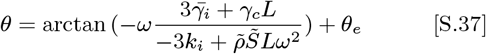

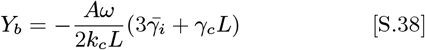

**Fig. S.17.**
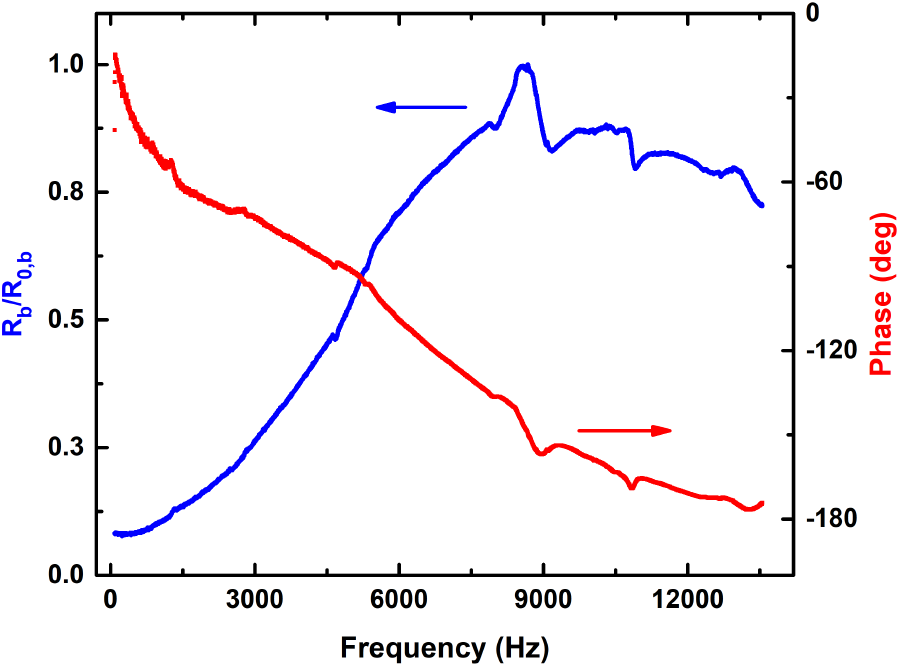
Frequency and phase response in 70% glycerol medium in commercial AFM. It is difficult to find truly off-resonance regime in these curves.

**Fig. S.18.**
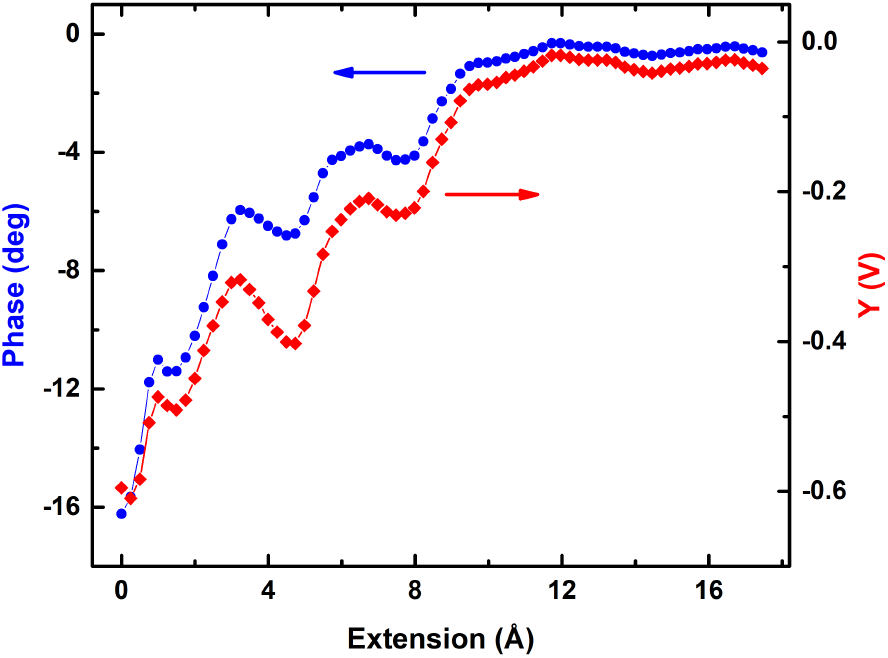
The raw data from which dissipation coefficient in Fig.9 is computed. The measurements are performed with interferometer based AFM. The phase for the cantilever displacement is zero when immersed in liquid and is far from the substrate. The data shows both the phase and Y-component of the displacement amplitude. They are used to calculate dissipation coefficient using point-mass and the continuous rectangular beam models respectively.

### Minimum detectable dissipation

The minimum detectable dissipation coefficient using our interferometer based AFM can be estimated by measuring the noise in Y-signal when the cantilever is not engaged with the substrate and is oscillating freely in the liquid. The displacement of the cantilever does not have any phase lag with respect to drive due to hydrodynamics alone. It means that the minimum detectable dissipation coefficient is related the minimum detectable Y-component. At the measurement bandwidth, dictated by the lock-in parameters, we looked at the broad-band noise of the Y-signal when the cantilever is free. The standard deviation of the this signal is then the minimum detectable Y-component of the amplitude. It turned out that the number is of the order of 5 pm. We arrive at the same number by recording and analysing the power spectral density of the cantilever displacement. From the noise floor and the lock-in time constant we estimated the noise in the cantilever displacement. Using this value of the noise in Y-component and Eq. 18, we obtain the minimum detectable dissipation coefficient to be 5 × 10^−7^*kg/s*. This is the upper bound on the dissipation coefficient of the unfolded single protein molecules.

